# High-Throughput Spatial Proteomics at Cellular and Subcellular Resolution via a Hanging Droplet Workflow

**DOI:** 10.1101/2025.11.06.686925

**Authors:** Hao Chen, Ulises H. Guzman, Jesper V. Olsen

**Affiliations:** Novo Nordisk Foundation Center for Protein Research, Department of Cellular and Molecular Medicine, Faculty of Health and Medical Sciences, University of Copenhagen, Copenhagen, Denmark

## Abstract

Formalin-fixed paraffin-embedded (FFPE) samples are essential for clinical research and spatial proteomics (SP) but are technically challenging to analyze by liquid-chromatography coupled mass-spectrometry (LC-MS) due to losses during protein extraction, evaporation during *de*crosslinking and low-throughput sample preparation methods. Here, we present a streamlined micro-FFPE proteomics sample preparation protocol for laser-capture micro-dissection (LMD) that enables scalable and high-throughput processing in 96-well format, completing the entire process from tissue lysis to tryptic peptide mixtures on LC-MS-ready Evotips in just 2-hours. Our method integrates whole-slide imaging, LMD, and ultra-sensitive narrow-window data-independent acquisition (nDIA)-MS and facilitates enhanced lysis, *de*crosslinking, and proteolytic digestion directly within Teflon-based EVO96 chips. By utilizing nano-wells and the hanging droplet concept, our approach significantly reduces surface adsorption, thereby improving processing efficiency for both higher- and very low-input FFPE samples. This enables quantification of >5,500 protein groups (PGs) by nDIA-MS from 50,000 µm² (∼250 cells) with robust performance maintained down to 1000 µm^2^ (∼5 cells) with ∼2,500 PGs identified. Application to a panel of breast cancer (BC) tissues precisely discerned BC subtypes and their tumor micro-environment (TME) based on differential MS-based protein abundance of biomarkers including TP53, HER2 and PGR. Individually excised subcellular regions (∼60 µm²) yielded ∼1,500 PGs, demonstrating reliable micro-scale spatial proteomics profiling approaching near-organelle resolution.

**Teaser:** Streamlined micro-FFPE protocol enables scalable and high-throughput processing of LMD samples, from lysis to mass spectrometry in 2 hours.

## Introduction

Eukaryotic cells in tissues and organs are embedded in highly organized tissue architectures that facilitates specialized biological functions^1^. The identity and function of cells are ultimately defined by their individual proteomes, however, it is equally important to consider the spatial distribution of these proteins within tissues and subcellular compartments. This spatial context is crucial for interpreting how proximity to neighboring cells, access to signaling cues, and positioning within multicellular networks influence cellular function. The emerging field of spatial proteomics addresses this need by capturing protein abundance differences within intact tissue architecture, moving beyond the averaged proteome profiles obtained from bulk and dissociated single cell analysis. For example, Deep Visual Proteomics (DVP) is a cutting-edge technology that integrates high-resolution imaging, artificial intelligence-driven cellular phenotype analysis, laser-capture microdissection, and ultra-sensitive mass spectrometry to map thousands of proteins at single-cell resolution while preserving spatial context. By linking proteomic information to precise morphological features at cellular and subcellular resolution, spatial proteomics enables the identification of cellular heterogeneity and dynamic intercellular communication that would otherwise remain hidden in complex environments, such as the tumor micro-environment (TME) or in intricate neuronal signaling networks^2,3^.

A uniquely enabling resource for spatial proteomics is the vast number of global archives of Formalin-Fixed Paraffin-Embedded (FFPE) human tissue blocks^4^. These worldwide archived samples, preserved as part of routine clinical practices, retain fine morphological structure and are often paired with well annotated clinical histories. These samples represent an unparalleled biobank of human biology across disease stages, tissue types and demographic backgrounds. The chemical crosslinking used for preservation is essential for maintaining tissue architecture and enabling precise pathological annotation and LMD, but it also makes protein extraction challenging^5^. Therefore, developing robust sample preparation methods to recover high-quality data from FFPE samples is critical for translating spatial proteomics into widespread clinical and translational research applications^6^. A central bottleneck in spatial proteomics workflows lies in reversing the formaldehyde-induced covalent methylene bridges between proteins, a process that requires high temperatures (typically >90 °C) to restore protein accessibility for enzymatic digestion^7^. While the high-temperature *de*crosslinking approach is effective for bulk samples, it poses a major challenge for micro-scale samples or thin tissue sections. The extremely small reaction volumes used in these preparations are highly prone to rapid evaporation, often occurring before *de*crosslinking is complete. As the sample dries, three critical problems arise. First, the tissue can be physically lost or damaged, directly reducing proteomics coverage. Second, premature drying interrupts the *de*crosslinking process and prevents efficient enzymatic digestion, leading to severely reduced peptide recovery and failure of the overall sample preparation workflow^8^. Thirdly, as the evaporation is difficult to control, it is often variable across FFPE samples leading to loss of sample preparation reproducibility. To mitigate these challenges, some approaches rely on the continuous replacement of buffer with in a heating system^9^. While effective, such approaches depend on specialized instrumentation, limiting their scalability and broader applicability. More recently, innovative “hanging droplet” strategies have been developed to mitigate evaporation in a simpler and more practical manner. For instance, a “hang-down” droplet configuration has been implemented to enhance *de*crosslinking and peptide generation from microscale samples by minimizing evaporation during heating^10^. Similarly, the Surfactant-assisted One-Pot voxel method (wcSOP) employs a suspended droplet on the tube cap to process laser-capture microdissected material, improving sample retention and proteomic depth^11^. However, despite their effectiveness in preventing evaporation and enhancing sensitivity, these approaches remain inherently low-throughput, as they require processing samples individually. Here, we introduce a one-pot, high-throughput spatial proteomics workflow capable of processing hundreds of LMD tissue regions in parallel. This method leverages of the EVO 96 chip to directly collect samples from a laser-capture microdissection microscope system, using a suspended droplet format to prevent evaporation during thermal *de*crosslinking and maintaining stable reaction volumes. This enables robust proteome profiling from extremely limited starting material, identifying ∼5,500 protein groups (PGs) from regions containing ∼ 250 cells and ∼ 2500 PGs from regions containing ∼5 cells using an 80 samples-per-day (SPD) LC-MS workflow. Moreover, our method achieves subcellular resolution, yielding 1,500 PG form subcellular structures as small as ∼60 µm² (e.g. isolated nuclei). Finally, we demonstrate the applicability of our method by profiling breast cancer tissues from four distinct molecular subtypes, accurately resolving tumor-and microenvironment-specific proteomic signatures.

## Results

### High-Throughput Scalable Workflow optimization

Spatial proteomics (SP) faces several technical challenges that can limit its effectiveness, particularly during sample preparation. One major issue is sample loss during protein extraction, which can adversely affect both the quantity and quality of the resulting data. Additionally, evaporation during the *de*crosslinking process threatens sample integrity, complicating subsequent analyses. Current low-throughput sample preparation methods often rely on specialized equipment, creating barriers to broader adoption. To improve the adoption of spatial proteomics, it is crucial to develop streamlined sample preparation techniques that minimize sample loss and evaporation while enhancing overall throughput. To achieve this goal, we have developed a formalin-fixed paraffin-embedded (FFPE) sample preparation protocol that integrates the hanging droplet concept^12,13^. This approach minimizes sample loss, enhances *de*crosslinking efficiency, and circumvents the need for specialized equipment during sample preparation, as the sample is directly excised from the FFPE samples into a reaction chamber formed by a droplet on Teflon-based capillaries of EVO96 proteoCHIP chip (Fig.1A). The hanging droplet concept utilizes droplets that remain suspended from the chip pillars when the EVO96 chip is placed face-down over a standard 96-well plate and heated to 95°C. Droplet stability is governed by the interplay of surface tension, gravity, and vapor pressure, particularly during *de*crosslinking and digestion. This configuration effectively suppresses evaporation, maintaining stable reaction volumes throughout the crosslinking process, with minimal volume loss (∼1 µL; Supplementary Fig. 1A). The protective effect is achieved through the establishment of a pressure differential during heating, which prevents droplet evaporation and ensures consistent sample processing.

**Fig.1.**
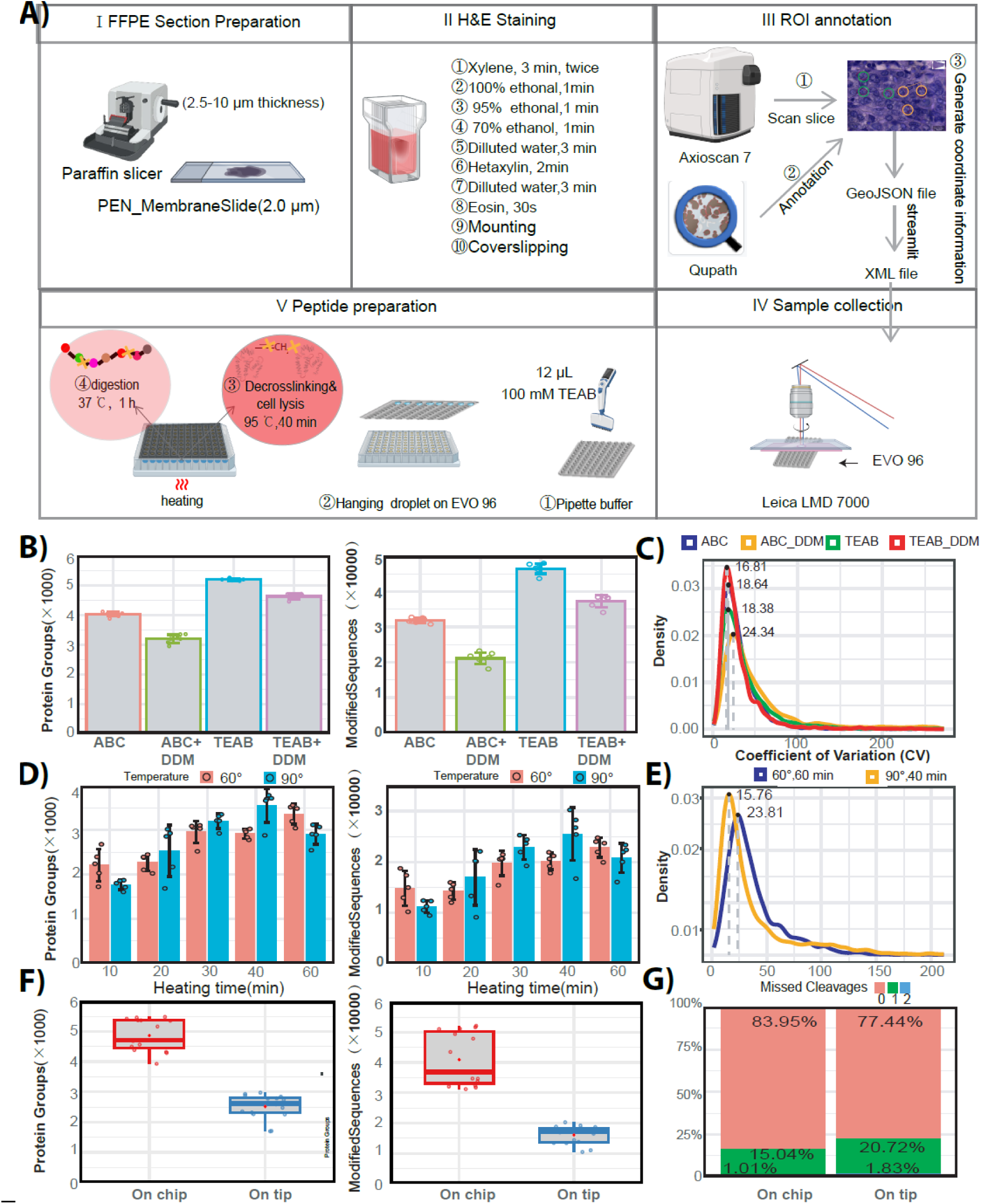
High-throughput hanging droplet workflow for micro-FFPE spatial proteomics. **(A)** Schematic overview of the integrated workflow combining whole-slide imaging, laser capture microdissection, and suspended droplet sample processing on EVO96 chips. **(B)** Comparison of Protein Groups and Modified Sequences identified using Ammonium Bicarbonate (ABC) versus Triethylammonium Bicarbonate (TEAB) digestion Buffers, with or without DDM. **(C)** Distribution of Coefficients of Variation (CV) across different buffer systems used in the digestion process. **(D)** Optimization of decrosslinking temperature (Blue: 90 °C, Red: 60 °C). **(E)** Distribution of Coefficients of Variation (CV) across different buffer systems used in the digestion process **(F)** Benchmark comparison between On-Chip (Red) vs On-tip (Blue) digestion. **(G)** Percentage of peptides containing up to two missed cleavage sites between On-Chip vs On-tip digestion.

To identify the optimal buffer conditions for the Lys-C and trypsin digestion process occurring in hang-down droplets, we compared two commonly used buffering systems: Ammonium Bicarbonate Buffer (ABC) and Triethyl ammonium Bicarbonate (TEAB), each tested with and without the addition of n-Dodecyl β-D-maltoside (DDM) on LMD-excised micro-FFPE breast cancer sections of 20,000 µm² with a thickness of 10 µm. Addition of the MS-compatible surfactant DDM was tested as it is known to prevent proteins from adsorbing to plastic surfaces, which is a major challenge in analyzing tiny samples like single cells and LMD-excised microFFPE material. Notably, in both buffer systems, omitting DDM resulted in higher numbers of identified PGs and modified peptide sequences. Among the tested buffers, TEAB provided the best performance, resulting in greater protein and peptide recovery as well as improved reproducibility, as reflected by lower coefficients of variation (CVs) (Fig.1B, C). The same trend was observed during the *de*crosslinking step (Supplementary Fig. 1B,C). This effect likely reflects the competing roles of the detergent: while DDM can reduce sample loss by limiting surface adsorption, it also alters the droplet’s surface tension and thermal behavior, leading to increased evaporation and reduced droplet stability. To further optimize the workflow, we systematically varied the proteolytic digestion time (15–90 min), the *de*crosslinking time (10–60 min) and temperature (60 and 90 °C). We found that 60 min digestion produced the highest number of identified protein groups and modified peptide sequences, while maintaining a similar proportion of missed cleavages compared to a 90 min digestion (Supplementary Fig. 1D,E). For the *de*crosslinking step, a 40-minute incubation yielded the greatest number of identified protein groups and modified peptides. Additionally, adjusting the *de*crosslinking temperature to 90 °C improved both protein and peptide recovery while also enhancing reproducibility across replicates (Fig. 1D, E). Together, these optimized conditions enable the preparation of LC-MS-ready samples in a 96-well format in approximately 2 hours. Finally, we compared our workflow to on-tip digestion^14^. Importantly, we found that the hanging droplet approach enables identification of twice as many protein groups and more than twice as many modified peptide sequences compared to on-tip digestion, along with a lower proportion of missed-cleaved tryptic peptides (Fig. 1F, G).

### Deep Single-Shot FFPE Proteome Coverage across Sample Loads with short gradients

To assess our FFPE sample preparation approach, we employed the 80 samples-per-day (SPD) whisper Zoom liquid chromatography (LC) gradient from the Evosep One system, coupled through an electrospray source with the Orbitrap Astral mass spectrometer. This setup utilized the narrow-window data-independent acquisition (nDIA) method, which has demonstrated the sensitivity required for analyzing sample amounts at the single-cell level^15,16^. Additionally, to evaluate the proteome coverage of FFPE samples, we measured the area of FFPE breast cancer samples, which ranged from 400 µm² to 50,000 µm², with a 10 µm thickness, corresponding to approximately 5 to 250 cells (Supplementary Fig.2A). As expected, the number of identified protein groups and modified peptide sequences scaled with input amount, yielding approximately ∼5,500 PGs and ∼50,000 modified peptides from the highest input (∼250 single cells), and ∼2,500 PG and ∼13,000 modified peptides at the lowest input (∼5 single cells), representing 42% of the proteins detected at the highest load (Fig.2 A, Supplementary Fig. 2B). Notably, we observed that smaller sampled regions exhibited greater biological heterogeneity, resulting in reduced reproducibility, consistent with our measurements (Fig. 2B). Nonetheless, low-input samples showed ∼85% overlap in identified protein groups and a Pearson correlation coefficients of ∼0.9 in their quantitative profiles, indicating high reproducibility and consistency even at minimal input levels (Fig. 2 C, D). While larger samples exhibited higher overall protein intensity (Fig.2E, F), the gene ontology (GO) term enrichment analysis demonstrated that, despite slightly lower identifications in small-input regions, the core functional process remained consistent (Supplementary Fig. 2C).

**Fig. 2.**
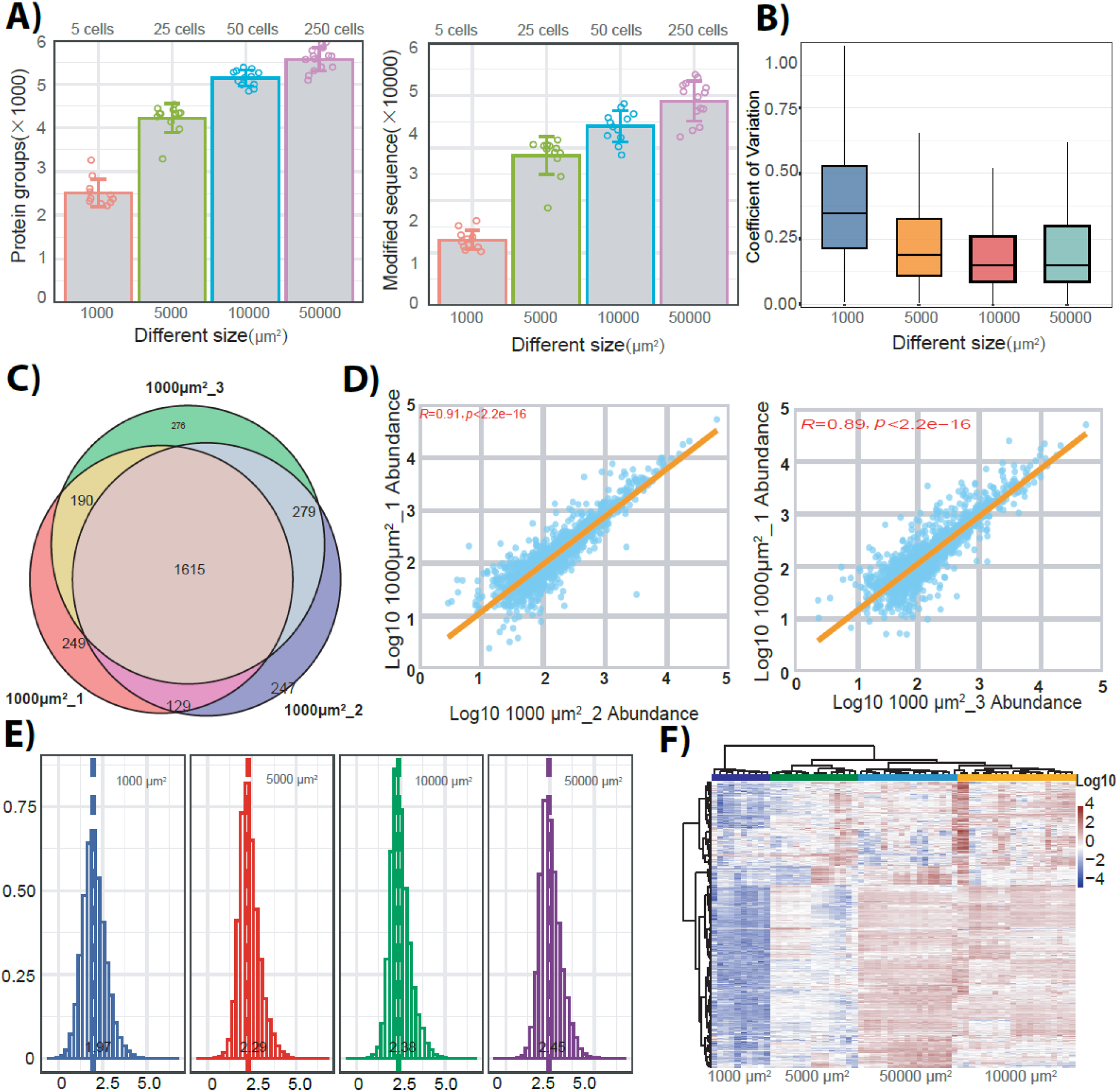
Single-shot proteome coverage across microdissected FFPE inputs. **(A)** Protein group and modified sequence identifications across different microdissected region size inputs, ranging from ∼50,000 μm² (≈250 cells) to ∼1,000 μm² (≈5 cells). **(B)** Box plots denoting the coefficient of variation (CV) across different microdissected region size inputs. All box plots include the median line, the box denotes the interquartile range (IQR), whiskers denote the rest of the data distribution and outliers are denoted by points greater than ±1.5 × IQR. **(C)** Venn diagram showing protein group identifications overlap between biological replicates. (n =3, 1000 μm^2^ input)**. (D)** Scatter plot showing the Pearson correlation across biological replicates. **(E)** Protein abundance distributions across different microdissected region size inputs **(F)** Heatmap displaying unsupervised hierarchical clustering across different microdissected region size inputs.

### Spatial characterization of the tumor microenvironment within triple-negative breast cancer (TNBC) tissue samples

While bulk proteomics of cancer-invaded tissues can provide deep insights into the state of the disease, this method has the limitation of averaging the protein signals from all the different cells involved in the disease process^17^. In contrast, single-cell proteomics (SCP) based on single cell suspensions can detect molecular signals from individual cells but loses spatial information^18,19^, which restricts the investigation of the complex interactions between malignant and non-malignant cells. Conversely, spatial proteomics allows for the capture of molecular signals at single-cell resolution while preserving spatial context^20,21^. To test this, we collected cancer and stroma cell regions of 20,000 µm^2^ from 10 µm thick FFPE triple-negative cancer breast samples (Supplementary Fig. 3A). Reassuringly, our method enables us to identify approximately 4,500 and 5,500 protein groups, alongside around ∼33,000 and ∼50,000 modified sequences from stromal and cancerous regions, respectively, with a good overlap between the samples (Fig. 3A, B). Principal component analysis (PCA) revealed that spatial localization of the sampled regions was the primary source of variation in the samples (Fig. 3C). The heatmap analysis revealed distinct clustering patterns based on the sample regions and protein intensity (Fig. 3D). The differential protein abundance analysis between stromal and cancerous regions identified 398 protein groups upregulated in cancerous regions and 407 protein groups upregulated in stromal regions, respectively (Fig. 3E). Among the proteins upregulated in the tumor regions, we identified key cancer-related proteins such as KRT8, FTH1, TOP2A, CDKN2A, which are recognized hallmarks of triple-negative breast cancer (TNBC)^22–25^. In fact, it has been reported that KRT8 plays a role as a luminal marker overexpressed in invasive epithelia^26^, whereas TOP2A, an enzyme essential for DNA replication and cancer cell proliferation, serves as a direct or indirect target of several cytotoxic anti-cancer agents^22^. Moreover, the FTH1 has been identified as a prognostic marker for triple negative breast cancer^24^. In contrast, the stromal regions showed a distinct molecular signature, with proteins such as COL1A1, COLA2, DCN and LUM upregulated in the stroma. In the stromal microenvironment, fibrillar collagens are known to accumulate in breast cancer and often appear as poorly defined spiculated masses in mammographic imaging^27^. Their abundance has been associated with multiple clinical pathological features, including tumor progression and patient outcome^28^. Additionally, the proteoglycans DCN and POSTN exhibit context-dependent roles, frequently functioning as endogenous regulators that modulate growth factor signaling and collagen organization, thereby helping to restrain tumor progression^29,30^. Consistent with these observations, gene ontology (GO) and KEGG pathway analyses revealed that stromal regions were enriched in biological processes such as extracellular matrix organization, immune response, B cell-mediated immunity, and wound healing (Supplementary Fig. 3B), indicating active fibrotic and immune processes. In contrast, tumor regions were enriched for process including RNA splicing, ribonucleoprotein complex biogenesis, and mRNA metabolic pathways (Supplementary Fig. 3C), supporting enhanced cellular proliferation and invasive potential. The compartment-specific protein and pathway signatures identified here are in strong agreement with known biological features of tumor and stromal microenvironments, demonstrating the robustness of our workflow in capturing molecular characteristics of the tumor microenvironment.

**Fig. 3.**
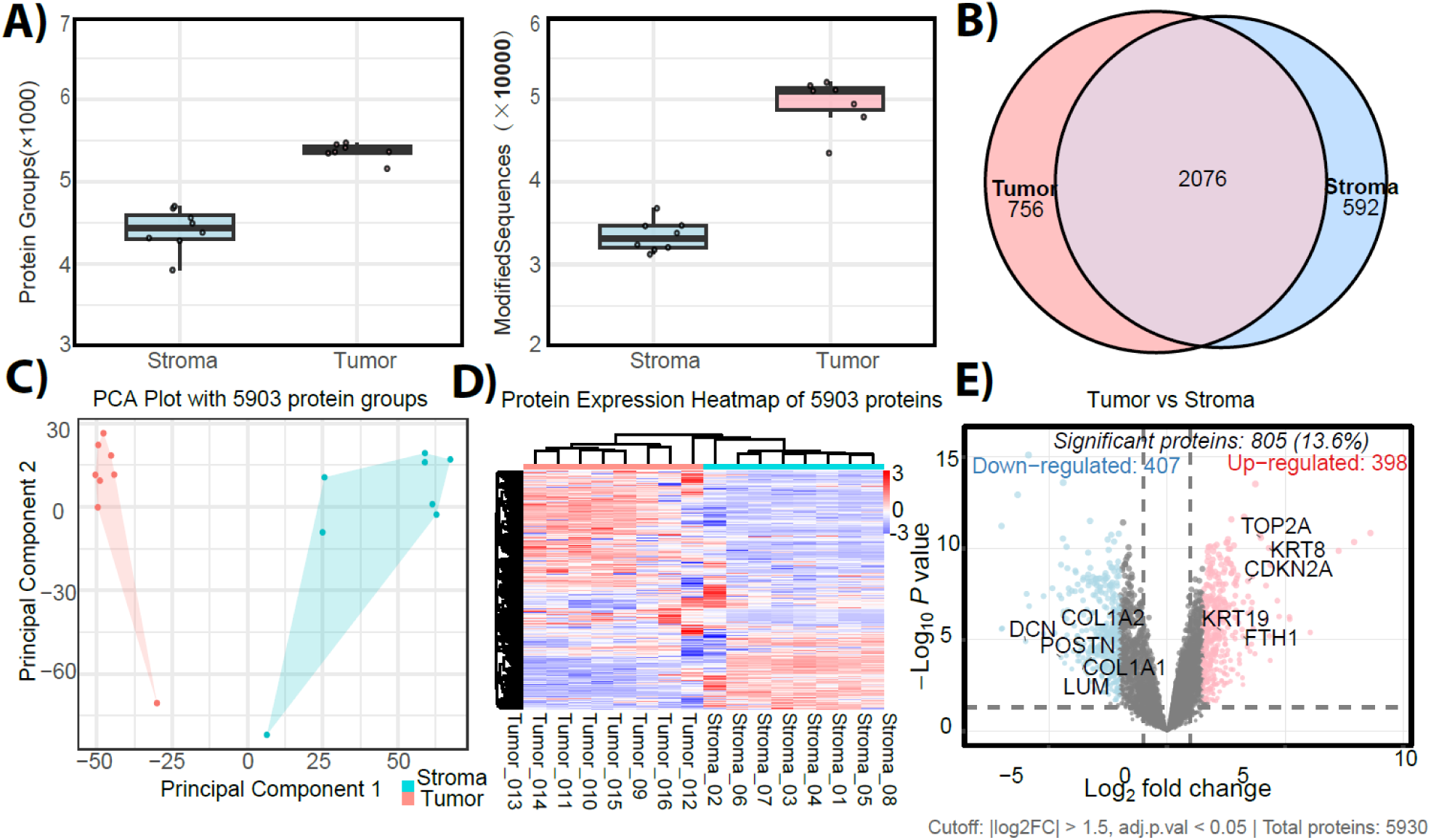
Spatial proteomics resolves tumor and stromal proteomic microenvironment in TNBC. **(A)** Protein group and modified peptide identifications from tumor and stromal regions collected from FFPE TNBC tissue. **(B)** Venn diagram showing protein group identifications overlap between cancer and stromal microdissected regions. **(C)** Principal Component analysis (PCA) depicting differences between tumor and stromal samples based on spatial origin. **(D)** Heatmap displaying distinct clustering of tumor vs. stromal proteome signatures. **(E)** Differential expression profiling of the Cancer and Stromal microdissected regions in a volcano plot. Significant regulated proteins at 1% false discovery rate (FDR) are delimited by dashed lines, respectively (FDR controlled, two-sided *t* test, randomizations = 250, s0 = 0.1).

### Spatial characterization of the tumor microenvironment in triple-negative breast cancer (TNBC) at subcellular resolution

While spatial proteomics has primarily focused on tissue-level protein mapping, an emerging frontier is subcellular proteomics, which aims to resolve protein distribution at organelle-scale resolution^31^. This approach is essential for understanding molecular function in situ. Yet conventional approaches rely on organelle enrichment from bulk tissue, obscuring cellular heterogeneity. Therefore, we tested our optimized micro-FFPE workflow for high-sensitivity subcellular spatial-proteomics profiling, by collecting micro-dissected ∼60 µm² regions from 5 µm-thick HER(–) ER(+) PR(+), P53(+) sections, targeting distinct tumor and stromal areas. Although the resolution of laser-capture microdissection (LCM) does not allow isolation of single, pure nuclei, the sampled regions were enriched for single nuclei, enabling robust proteomic profiling across tissue compartments (Supplementary Fig. 4A). Proteome profiling of individual nuclear-enriched regions yielded slightly higher identifications on tumor than stroma (∼1,500 vs ∼1,400 PGs; ∼8,500 vs 7,500 modified peptides; Fig. 4A), although the two proteomes were largely overlapping (Fig.4 B). PCA further resolved the samples into distinct tumor and stromal nuclear states (Fig.4 C). The differential expression analysis of the nuclear-enriched regions identified 115 proteins upregulated in tumor and 90 PGs in stroma. Tumor nuclei were enriched for cancer- and nucleus-associated proteins, including TP53, BRCC3, ELAVL1, and PSMC4, whereas stromal nuclei showed elevated levels of proteins linked to extracellular matrix organization, fibrosis, and immune regulation, such as COL2A1, COL12A1, IGLC3, and IGHV3 (Fig. 4D). These patterns are consistent with the known biological distinctions between tumor and stroma and recapitulate trends previously observed from larger-scale sampling of similar tissue regions. Although estrogen receptor (ER) and progesterone receptor (PGR) were not detected, we identified and quantified clinically relevant cancer markers, including ERBB2 (Receptor tyrosine-protein kinase erbB-2) also known as HER2, and TP53, with the latter showing a significant difference between tumor and stromal nuclei (Fig. 4E). To further assess the sensitivity of our workflow at the subcellular level, we collected 1, 10 and 50 sub-cellular enriched nuclei samples (5 replicates per conditions). As expected, base peak intensity increased proportionally with the sample input (Supplementary Fig. 4B), and the number of identified protein groups ranged from ∼1,200 to ∼1,800, with ∼7,000 to ∼ 13,000 unique peptides detected (Fig. 4F). Unsupervised hierarchical clustering revealed sample grouping based on input material (Fig. 4G), and most nucleosome subunit proteins were detected even at the single-enriched nuclei level (Fig. 4H). Similarly, the Pearson correlation analysis between samples showed the same trend (Supplementary Fig. 4C). GO analysis of the top 20 PGs detected showed strong enrichment for nuclear-related cellular components (Supplementary Fig 4D). Altogether, these results demonstrate that our optimized workflow can detect quantitative differences between subcellular structures from distinct histological regions while preserving biologically relevant information.

**Fig. 4.**
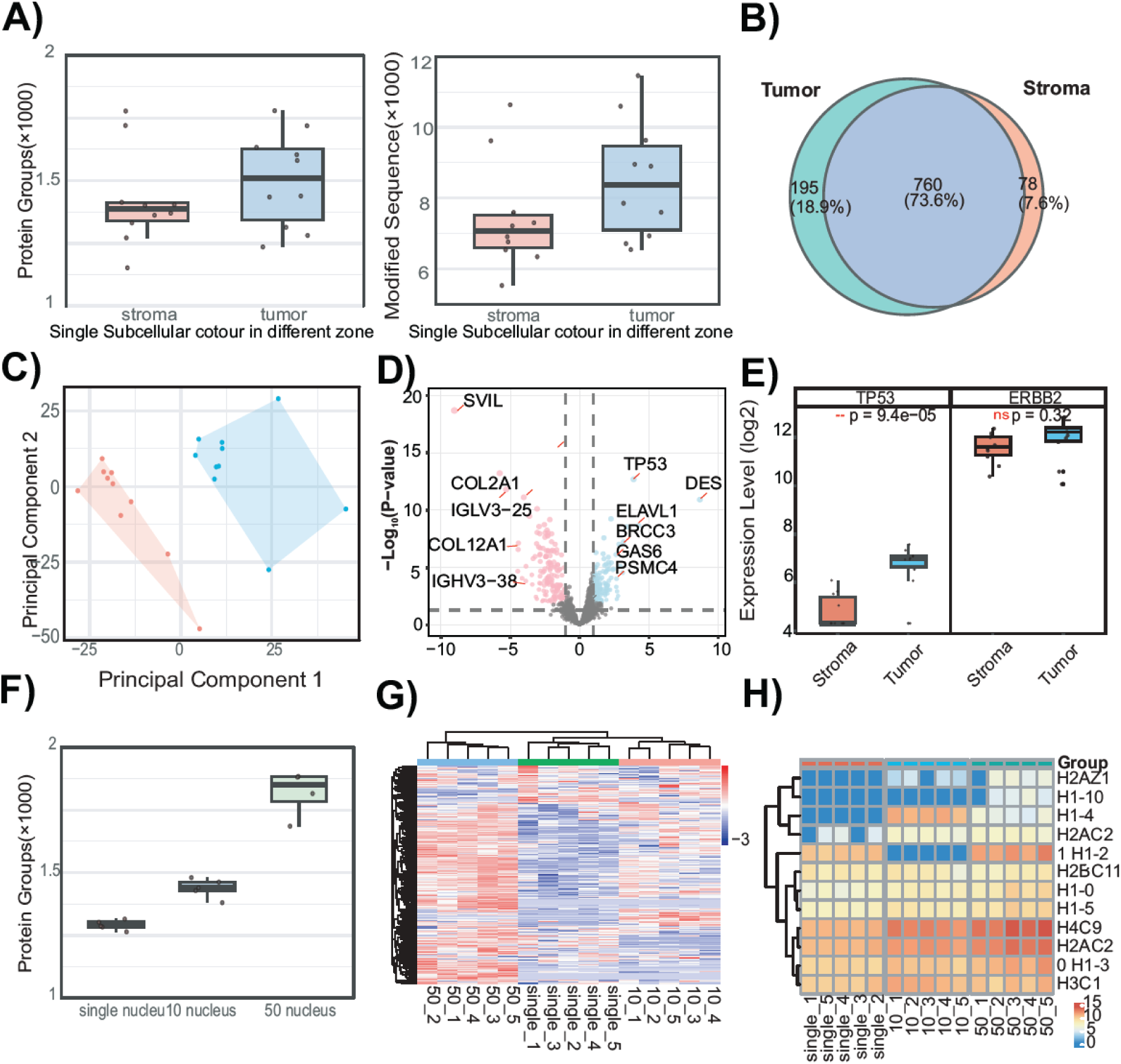
Subcellular-resolution spatial proteomics of nuclear-enriched micro-region. **(A)** Box plots depicting the proteome depth (Protein Groups and Modified sequences) from nuclear-enriched regions (∼60 μm²). **(B)** Venn diagram showing protein group identifications overlap between nuclear proteomes derived from distinct regions (Stroma and Cancer). **(C)** Principal Component analysis (PCA) depicting difference between tumor vs. stromal nuclear states. **(D)** Differential protein expression profiling enriches tumor and cancer nuclei in a volcano plot. Significant regulated proteins at 1% false discovery rate (FDR) are delimited by dashed lines, respectively (FDR controlled, two-sided *t* test, randomizations = 250, s0 = 0.1). **(E)** Boxplots depicting clinically relevant cancer markers (ERBB2 and TP53) at subcellular scale. **(F)** Boxplot displaying the protein group and peptide identifications from 1 to 50 nuclei inputs. **(G)** Heatmap showing clustering groups samples by input level. **(E)** Heatmap displaying the nucleosome coverage across distinct input amounts (n=5). All box plots include the median line, the box denotes the interquartile range (IQR), whiskers denote the rest of the data distribution and outliers are denoted by points greater than ±1.5 × IQR.

### Spatial profiling of the tumor microenvironment in breast cancer subtypes

The four major breast cancer subtypes are Luminal-A, Luminal-B, HER2-positive, and triple-negative. These categories are determined by the tumor’s status regarding the hormone receptors - estrogen receptor (ER) and progesterone receptor (PR) - and the human epidermal growth factor receptor 2 (HER2). Each subtype has different growth rates, prognoses, and responses to treatment. To investigate subtype-specific features of the tumor microenvironment across the four major breast cancer subtypes: HER2-positive (ER⁻/PR⁻/HER2⁺), Luminal-B (ER⁺/PR⁺/HER2⁺), Luminal-A (ER⁺/PR⁺/HER2⁻), and triple-negative (ER⁻/PR⁻/HER2⁻), we microdissected ∼4,000 μm² and 5 μm thickness regions (∼10 cells) from paired tumor and stromal compartments and analyzed them using our optimized spatial-proteomics workflow. Hierarchical clustering based on proteins significantly differing across subtypes (two-way ANOVA; FDR 0.05) revealed pronounced molecular divergence between regions (Supplementary Fig. 5A). Within tumor regions, the ER⁻/PR⁻/HER2⁻ (triple-negative) subtype showed strong enrichment for immune-related proteins, consistent with previous reports of increased immune infiltration and antigen-presenting cell engagement in triple-negative breast cancer (TNBC)^32,33^. In contrast, the HER2 and/or hormone receptor positive subtypes showed enrichment for proteins associated with translational machinery, aligning with luminal and HER2-driven proliferative programs^34^. Conversely, stromal regions exhibited subtype-specific extracellular matrix (ECM) remodeling signatures: ER⁻/PR⁻/HER2⁺, ER⁺/PR⁺/HER2⁺, and ER⁺/PR⁺/HER2⁻ stroma were enriched for collagen- and fibronectin-associated matrix organization, in agreement with known fibroblast-mediated ECM remodeling in hormone receptor– positive and HER2⁺ tumors^35,36^. In contrast, ER⁻/PR⁻/HER2⁻ stroma showed greater enrichment of translational and metabolic pathways, suggesting a distinct stromal activation state characteristic of TNBC-associated microenvironments^37^. To resolve molecular signatures across cancer subtypes and regions, we performed correlation analysis across all identified protein groups, irrespective of the cancer subtype, revealing co-regulated molecular modules. Seven distinct signatures were observed in tumor regions and four in stromal regions (Fig. 5A). Examination of tumor-associated modules highlighted subtype-specific differences; for example, the metastasis-associated cluster comprising proteins with significant subtype-dependent variation, showed upregulation of S100P and EVPL in the ER⁻/PR⁻/HER2⁻ (triple-negative) subtype, consistent with previous reports linking S100P to TNBC metastasis^38^(Supplementary Fig. 6A). Within the breast cancer-associated cluster, as expected, PR was detected in ER⁺/PR⁺/HER2⁺ and ER⁺/PR⁺/HER2⁻ subtypes ^38^(Supplementary Fig. 6B). Notably, the immune system-associated module exhibited upregulation of proteins including SERPINA1, SERPINB1, and LRG1; the latter is a glycoprotein known to promote angiogenesis and tumor progression, and its elevated expression in breast cancer has been associated with lymphatic metastasis and poor patient survival^39^(Supplementary Fig. 6C). In contrast, the metastasis-associated module derived from the stromal compartment showed the upregulation of proteasome subunits, including PSMA1, PSMB1 and PSMB2 across the majority of breast cancer subtypes analyzed. Notably, PSMA1 has been previously implicated in breast tumor progression, underscoring its potential role in supporting malignant growth^40^. More broadly, coordinated expression of proteasome gene signatures has been associated with high tumor grade and aggressive phenotypes in breast cancer, highlighting the contribution of proteasomal regulation to tumor malignancy^41^(Supplementary Fig. 7A). The matrisome-associated cluster exhibited upregulation of collagen matrix proteins (Supplementary Fig. 7B), including COL1A1, COL1A2, and COL6A1, compared with triple-positive and triple-negative breast cancer subtypes, suggesting differential collagen matrix deposition between cancer subtypes^42,43^. Moreover, the triple-negative subtype exhibited marked activation of Rho-GTPase signaling (Supplementary Fig. 7C), alongside elevated expression of S100A11 and S100A4, proteins associated with the promotion of the breast cancer cell invasiveness. Finally, to assess compartment-specific differences, we compared tumor and stromal regions within each subtype. This analysis reliably recovered the main cell-type– specific markers characteristic of each subtype with ERBB2 more abundant in cancer cells than stroma in HER2-positive and luminal-B samples, whereas PGR was more abundant in cancer cells than stroma in luminal-A and luminal-B samples, respectively (Fig. 5B). Across most subtypes, we observed broad activation of the MYC pathway and epithelial-to-mesenchymal transition (EMT). In the triple-negative subtype, MYC pathway activation was detected predominantly in the stromal compartment, suggesting that stromal cells may contribute to the aggressive TNBC phenotype^44^. Alternatively, this signal could reflect cancer-cell infiltration into the stroma or paracrine induction of MYC signaling in stromal cells^45^.

**Fig. 5.**
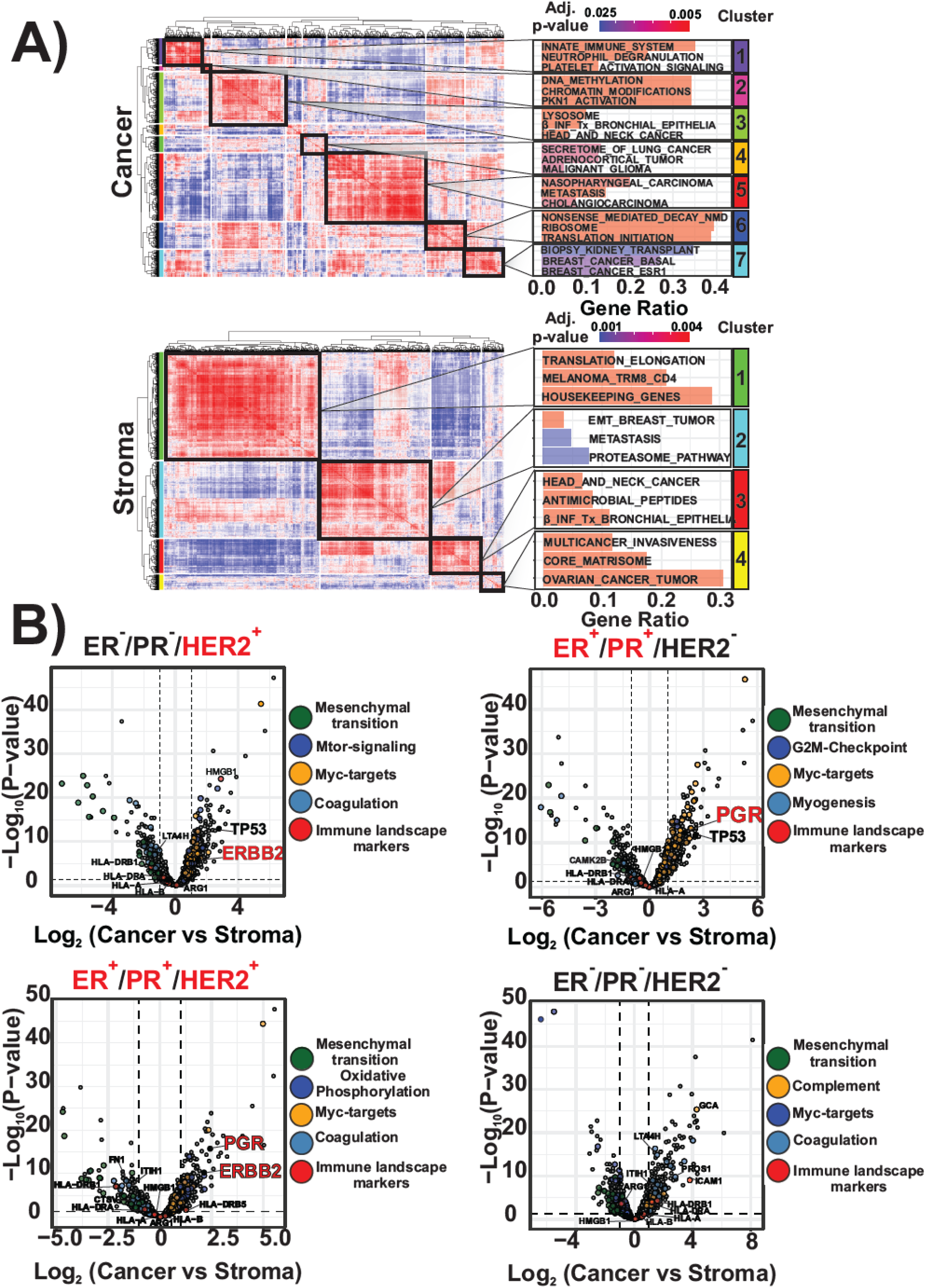
Spatial proteomic mapping of tumor and stromal compartments across breast cancer subtypes. **(A)** Pearson correlation analysis across all quantified protein groups identifies distinct co-regulated molecular modules within the tumor (upper panel) and stromal (lower panel) compartments among ER⁻/PR⁻/HER2⁺, ER⁺/PR⁺/HER2⁺, ER⁺/PR⁺/HER2⁻, and triple-negative (TNBC) subtypes. Functional enrichment by module and microdissected region is presented in the left panel. **B-D)** Differential protein expression profiling of tumor regions compared to the corresponding stromal regions by subtype nuclei in a volcano plot. Significant regulated proteins at 1% false discovery rate (FDR) are delimited by dashed lines, respectively (FDR controlled, two-sided *t* test, randomizations = 250, s0 = 0.1

## Discussion

Spatial proteomics aims to characterize protein expression within the structural context of tissues, enabling insights into cell–cell communication, microenvironmental influences, and functional specialization of cell states that are not captured by bulk or dissociated single-cell proteomics^46^. However, widespread adoption of spatial proteomics has been constrained by practical challenges, particularly the efficient extraction and digestion of proteins from microscale tissue regions. To address this, we introduce a high-throughput workflow based on the hanging droplet concept that enables reproducible processing of up to 96 laser-microdissected samples in parallel, generating LC-MS–ready material in ∼2 hours. Our streamlined approach achieves robust proteome coverage from low-input samples, recovering ∼5,500 PGs from input equivalent to ∼250 cells and ∼2,500 PGs from input equivalent to ∼5 cells. Moreover, the workflow supports reproducible subcellular-scale profiling, recovering ∼1,500 protein groups from nuclear-enriched regions as small as ∼60 µm², and approaching near-organelle resolution while preserving cell-type–specific molecular signatures. The biological application of this workflow to triple-negative breast cancer (TNBC) tissues revealed robust compartment-specific proteomic programs distinguishing tumor from stromal regions. Tumor regions were enriched for RNA processing pathways, proliferative signaling networks, and canonical cancer-associated markers, whereas stromal regions exhibited signatures of extracellular matrix remodeling, immune activation, and fibrotic repair. Extending this analysis across multiple breast cancer subtypes uncovered distinct molecular programs within both tumor and stromal compartments, highlighting subtype-dependent differences in micro-environmental composition and signaling dynamics. Notably, our approach demonstrates the ability to detect biologically meaningful protein signatures from regions representing as small as ∼60 μm², bridging the sensitivity gap between single-cell proteomics and tissue- or subcellular-level spatial resolution, without the need for highly specialized instrumentation. In summary, our scalable and reproducible workflow enables ultrasensitive spatial proteomics profiling of FFPE tissues at micro-regional and subcellular resolutions, bridging the gap between single-cell proteomics and tissue-level analyses. This advancement holds significant potential for translational research and precision pathology.

## Materials and Methods

### Materials

Formalin-fixed paraffin-embedded (FFPE) human triple-negative breast cancer tissue slides, measuring 5 µm and 10 µm in thickness, embedded in a 2.0 um PEN membrane glass slides were obtained from VitroVivo Biotech. (VitroVivo Biotech LLC P.O. Box 83644 Gaithersburg, MD 20883) and processed as follows.

### H&E staining

Prior to analysis, FFPE slides were dewaxed in xylene for 3 minutes, repeated twice. After dewaxing, samples underwent a graded hydration protocol: 100% ethanol for 1 minute (twice), 95% ethanol for 1 minute, 70% ethanol for 1 minute, and finally distilled water for 3 minutes. After dewaxing, samples underwent a graded hydration protocol: 100% ethanol for 1 minute (twice), followed by 95% ethanol for 1 minute, 70% ethanol for 1 minute, and finally distilled water for 3 minutes. Subsequently, the slides were stained with hematoxylin for 1 minute. Excess hematoxylin was removed with distilled water before staining with Eosin for 10 seconds, after which any remaining stain was also washed off with distilled water. For directly capturing samples using the laser capture microdissection (LCM) system, stained slides were air-dried and stored at −20°C. For annotation and more accurate sample capture based on clear images, samples were sealed with an anti-fade fluorescence mounting medium and covered with a glass coverslip.

### Visualization and annotation

Images captured at 20X or 40X magnification using the Axioscan 7 microscope are imported into QuPath 0.5.1, where regions of interest (ROIs) are marked and annotated before being exported as a GeoJSON file. The web application “Steamlist(https://qupath-to-lmd-mdcberlin.streamlit.app/)” then converts the GeoJSON file into an XML file, which contains the coordinate information necessary for the Laser Microdissection (LMD) machine to identify the samples. For sample collection. A proteoCHIP EVO 96 (cellenONE X1 Neo) is used as a holder, requiring an adaptor to secure the EVO 96 chip within the LMD shelter. This adaptor was 3D printed, with the original design from Makhmut et al., 2024^47^.

#### Laser capture

Prior to laser capture, the coverslips and mounting medium should be removed by incubating the slides in 1x PBS at 37°C for 30 minutes. Subsequently, the sections should be air-dried for 30 minutes at 37°C. The Leica LMD 7000 microscope was utilized for the laser capture and collection of samples, following the guidelines outlined in the XML file. The configuration for the laser parameters is as follows:

For large size sample capture (over 5000 µm^2^)

- Microscope: 10x
- Power: 60
- Apeture: 10
- Speed: 8
- Middle Pulse Count: 4
- Final Pulse: 08
- Head Count: 69%
- Pulse Frequency: 2200

For single cell or single subcellular level sample capture (less than 400 µm^2^)

- Microscope: 63x
- Power: 46
- Apeture: 1
- Speed: 10
- Middle Pulse Count: 4
- Final Pulse: 0
- Head Count: 55%
- Pulse Frequency: 2600

Following the laser capture and collection process, verification performed by microscope was conducted to determine whether the samples had been successfully deposited into the wells of the chip.

### Sample preparation by hanging on droplet(on)

To initiate the de-crosslinking process, 12 µL of 25 mM ABC buffer is added to the wells of the EVO 96 chip, which is positioned on a standard 96-well plate with the pillars facing inward. The assembly is then heated in a heating block at 95 °C for 40 minutes. After the heating period, the EVO 96 chips are carefully removed from the 96-well plate and allowed to cool to room temperature for 5 minutes. Subsequently, to initiate the digestion process 2.5 µL of a 10 ng/µL trypsin and Lys-C mix is added. The EVO 96 chips are then placed back on the 96-well plate and incubated at 37 °C for 1 hour. After digestion, the reaction is quenched by adding 1 µL of 10% formic acid (FA) to each well. Subsequently, samples are purified on EVOTIPS (Evosep) following the standard Evosep protocol and stored on EVOTIPS at −4 °C until LC-MS/MS analysis. Alternatively, the samples on the EVO 96 chips are suspended above the 96-well plate and stored at −20 °C until needed.

### Sample preparation by On-tip digestion

To initiate the *de*crosslinking process, 12 µL of 25 mM ABC buffer is added to the wells of the EVO 96 chip, which is positioned on a standard 96-well plate with the pillars facing inward. The assembly is then heated in a heating block at 95 °C for 40 minutes. After the heating period, the EVO 96 chips are carefully removed from the 96-well plate and allowed to cool to room temperature for 5 minutes. Thereafter, 2.5 µL of a 10 ng/µL trypsin and Lys-C mix are added. The proteolytic digestion was performed in an incubator at 37 °C for 1 hour. The resulting peptide mixtures were desalted and concentrated on C18-reversed-phase material in Evotips following the vendor’s instructions. Briefly, to activate the dried Evotips, they were washed with 20 μl of Solvent B (0.1% FA in acetonitrile) and centrifuged at 800 g for 60 seconds. To condition the Evotips, they were soaked in 100 μl of 1-propanol until they turned pale white. After this, they were equilibrated and with 20 μl of Solvent A (0.1% FA in water) and centrifuged at 800 g for 60 seconds. The sample were transferred to the Evotips by centrifugation at 100 rpm for 20 seconds. the Evotips were washed with 20 µL of Solvent A and centrifuged at 800g for 60 seconds. Then 100 µL of Solvent A were transferred to the Evotips and centrifuged at 800g for 10 seconds. Water was added to the Evotip box to the level of the C18 resin in the Evotips, andthe Evotips were stored at −4 °C until analysis by LC-MS/MS.

### LC separation and Mass analysis

All LC-MS/MS analysis was performed on an Orbitrap Astral MS using Thermo Tune software (version 1.0.100.40), coupled with an Evosep One system (EvoSep Biosystems) operated via Chronos software (Chronos 4.10.0.0, Chronos plugin 1.0.25.4). FFPE-derived peptide mixtures were analyzed using the predefined 80SPD whisper zoom method (5.6-minute gradient), using a 5-cm commercial analytical column (Aurora Elite TS, IonOpticks), interfaced online with an EASY-Spray™ source. The Orbitrap Astral MS was operated in narrow-window data-independent acquisition (nDIA) mode with a full MS resolution setting of 240,000 and MS1 injection time restricted to 3 ms, covering a full scan range of 380–980 m/z. The full MS AGC was set to 500%. MS/MS scans. The nDIA isolation windows were set to 4 Th with a maximum ion injection time (IT) restricted to 6 ms. The MS/MS scanning range was set to 380 to 980 m/z, with a normalized collision energy (NCE) of 27%.

### Proteomics raw data analysis

Raw LC-MS files were analyzed in Spectronaut v19 (Biognosys) with a spectral library-free approach (directDIA +) using the human protein reference database (Uniprot 2024 release, 20,588 sequences) for all samples complemented with common contaminants (246 sequences). Note, as the protocol does not involve reduction and alkylation, database searches were performed with free cysteine sulfhydryls and hence cysteine carbamylation was not set as a fixed modification, whereas methionine oxidation and protein N-termini acetylation were set as variable modifications. Precursor filtering was set to perform based on Q-values, and cross run normalization was checked. Method evaluation mode was enaled for method optimization.

## Acknowledgements

We would like to thank Juanjuan Wang and Frederik Post from the Mann Group at CPR for help with initial training on the LCM microscope.

## Author contributions

H.C., U.H.G., and J.V.O. designed the proteomics experiments, H.C. and U.H.G. prepared and performed proteomics experiments and analyzed the resulting data. J.V.O. critically evaluated the results. H.C., U.H.G., and J.V.O. wrote the first draft of the paper. All authors read, edited and approved the final version of the paper.

## Inclusion and ethics statement

We are committed to promoting diversity and inclusion in science and ensuring that our research is conducted ethically and responsibly. Our study was designed and conducted in accordance with ethical principles and guidelines, including obtaining informed consent from all participants and complying with relevant regulations and laws.

## Supplementary Materials

**Supplementary Fig. 1.**
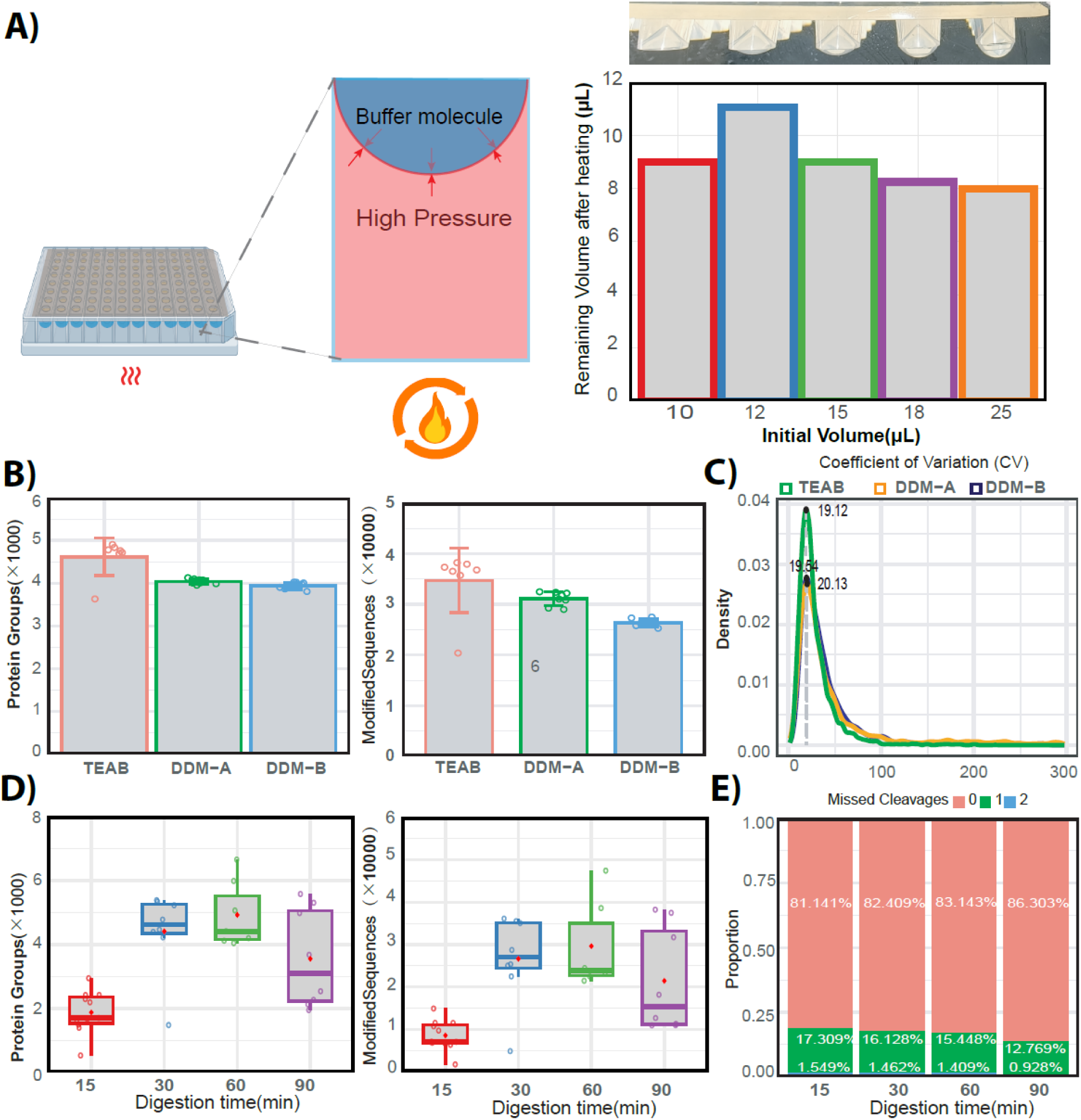
Conceptualization and optimization of the hanging droplet FFPE workflow. **(A)** Schematic illustration of the suspended droplet on the hanging droplet configuration on EVO96 chip pillars during decrosslinking (right panel). Reaction volume optimization during crosslinking conditions **(B)** Comparison of Ammonium Bicarbonate (ABC) and Triethylammonium Bicarbonate (TEAB) buffer systems adding DDM detergent before and after crosslinking process. **(C)** Distribution of Coefficients of Variation (CV) across different buffer systems adding DDM detergent before and after crosslinking process. **(D)** Boxplots illustrating protein group and modified peptide identifications across digestion time conditions. **(E)** Percentage of peptides containing up to two missed cleavage sites across distinct digestion time conditions. All box plots include the median line, the box denotes the interquartile range (IQR), whiskers denote the rest of the data distribution and outliers are denoted by points greater than ±1.5 × IQR.

**Supplementary Fig. 2.**
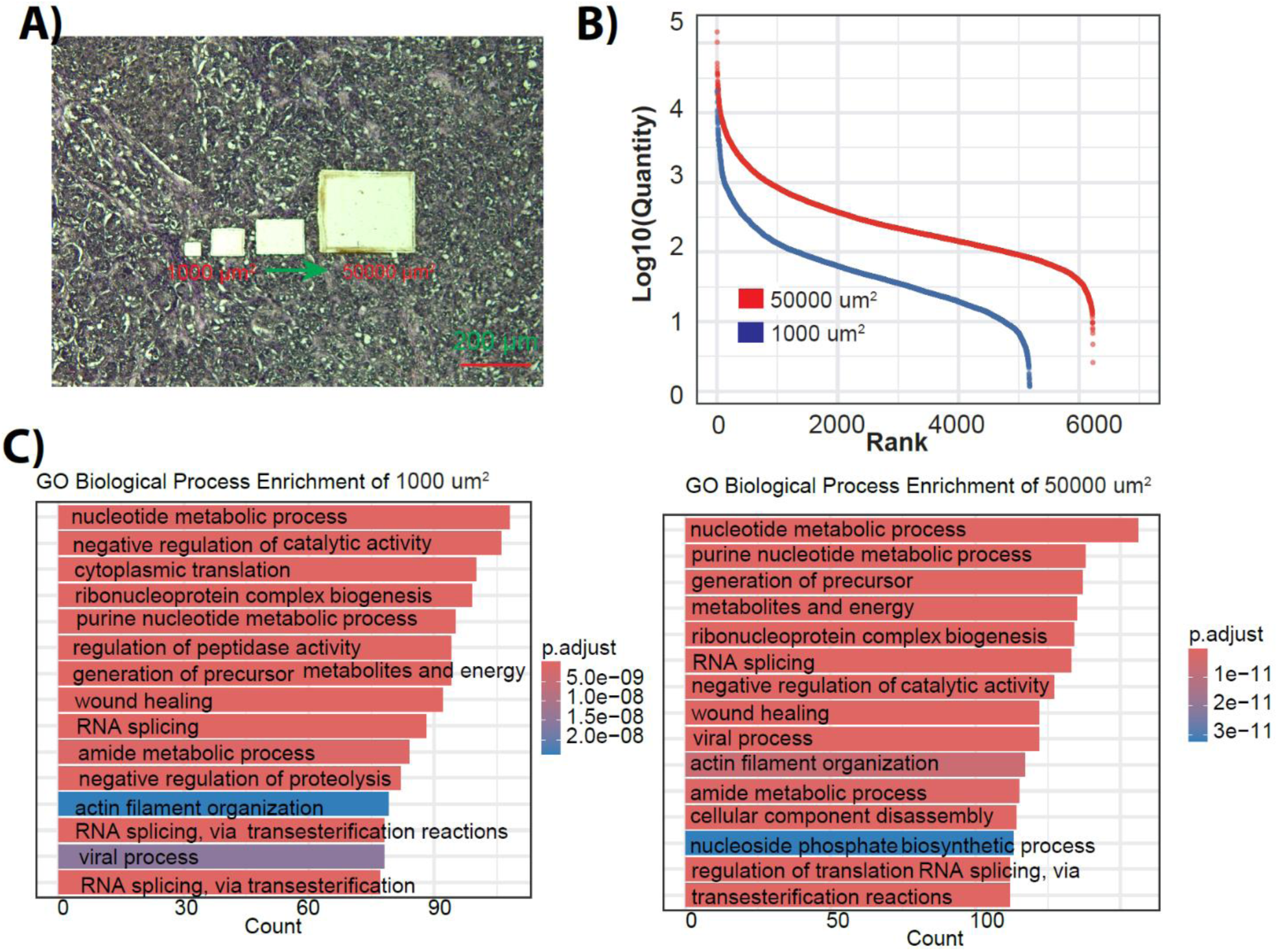
Proteome coverage and reproducibility across FFPE sample input amounts. **(A)** Microdissected region size representation ranging from ∼100 µm² to ∼50,000 µm² (∼5–250 cells). **(B)** Rank intensity plots showing the dynamic range across loading levels. **(C)** GO enrichment analysis across input amounts.

**Supplementary Fig. 3.**
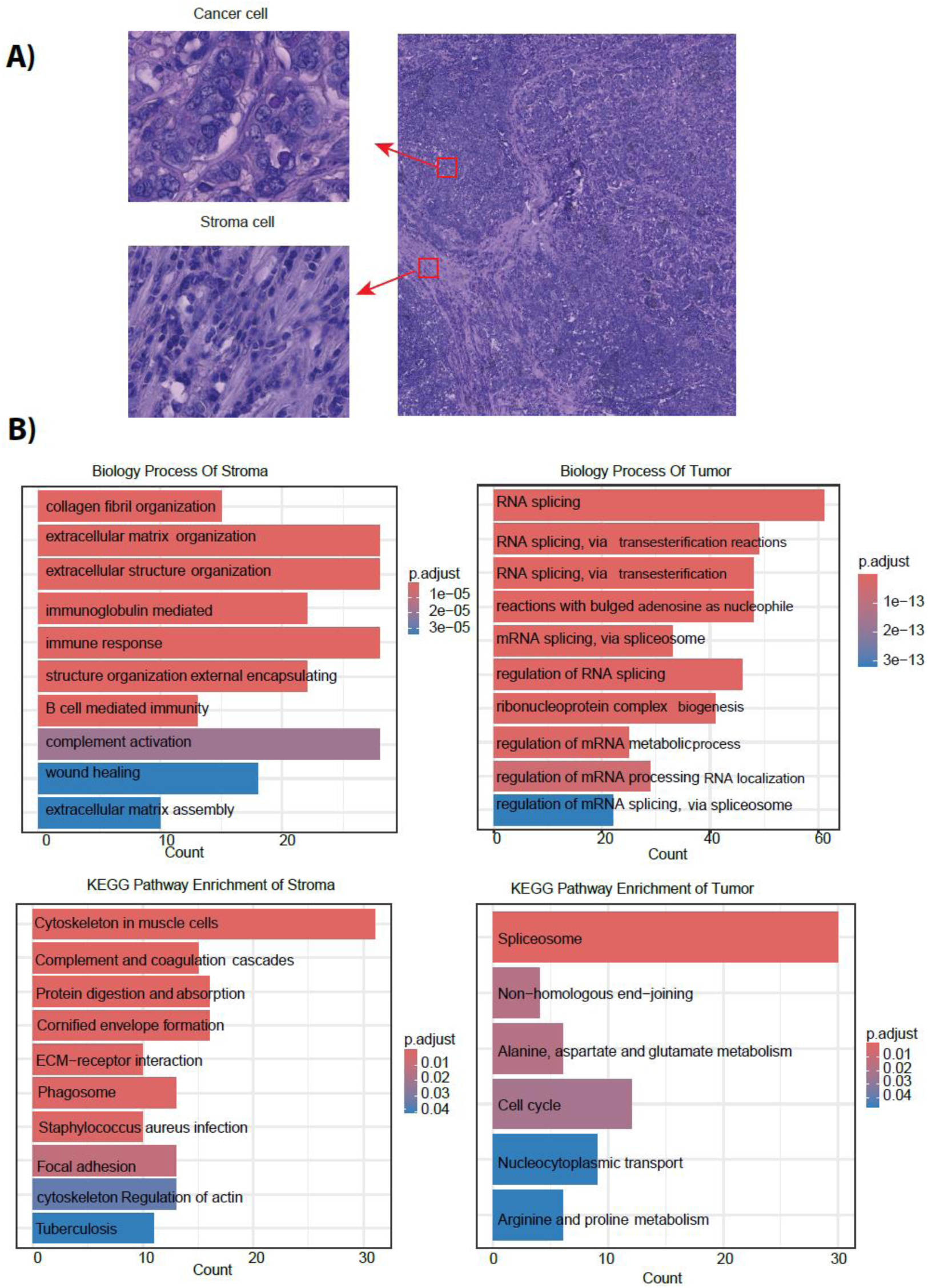
Spatial proteomic differentiation of tumor and stromal regions in TNBC. **(A)** Representative annotated tissue sections showing cancer and stromal microdissected regions. **(B)** Go enrichment-Biological process (upper panel) and KEGG pathway enrichment analysis of stromal regions (lower panel). (C) Go enrichment-Biological process (upper panel) and KEGG pathway enrichment analysis of tumor regions.

**Supplementary Fig. 4.**
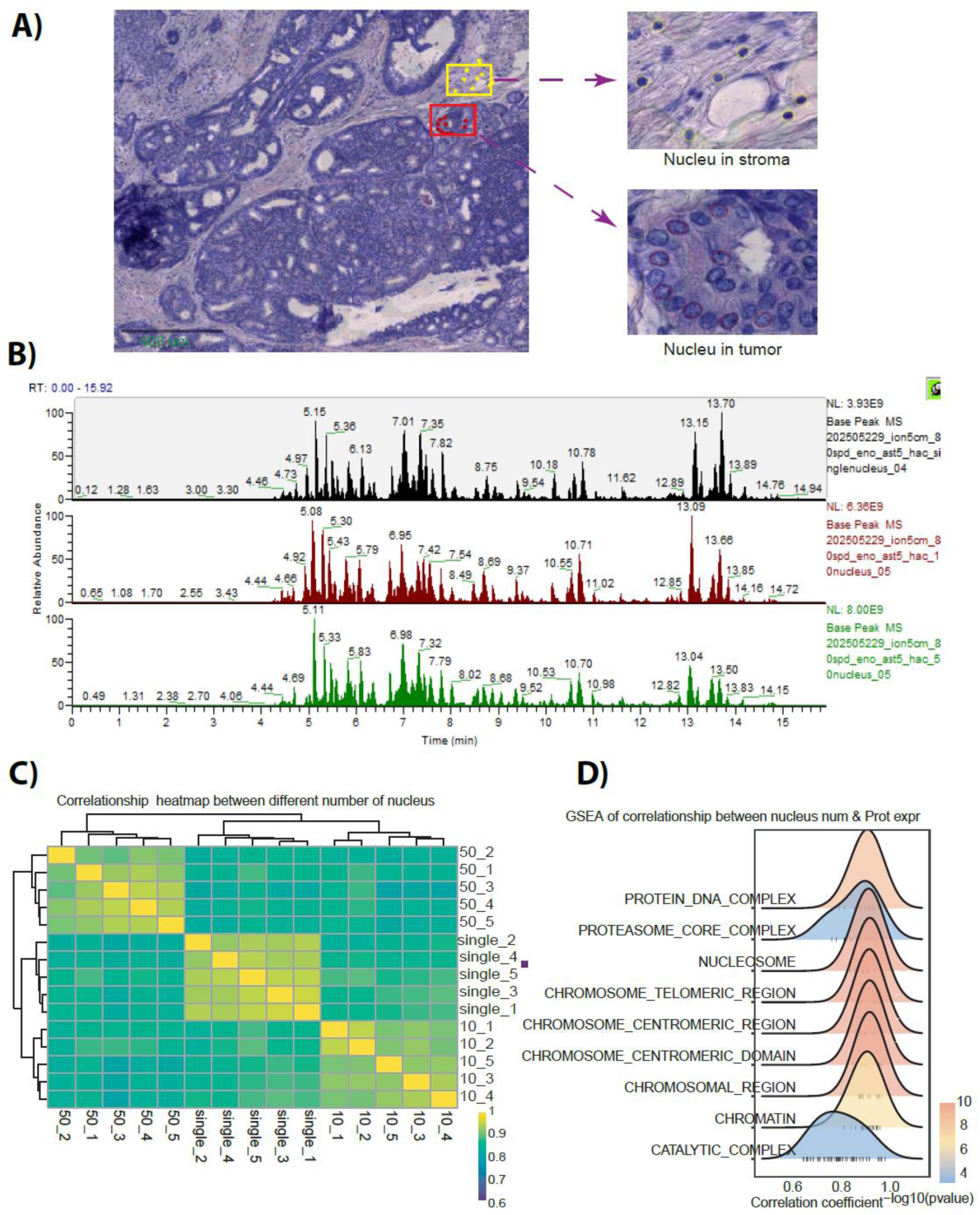
Subcellular spatial proteomics of nuclear-enriched micro-regions. **(A)** Microdissected nuclear-enriched regions (∼60 µm²) from tumor and stromal compartments. **(B)** Representative base peak chromatograms showing peptide signal scaling across input levels of 1, 10, and 50 nuclei. **C)** Pearson correlation and clustering analyses reflect consistent quantitative proteome profiling across biological replicates. **(D)** GESEA enrichment analysis of nuclear enriched microdissected samples.

**Supplementary Fig. 5.**
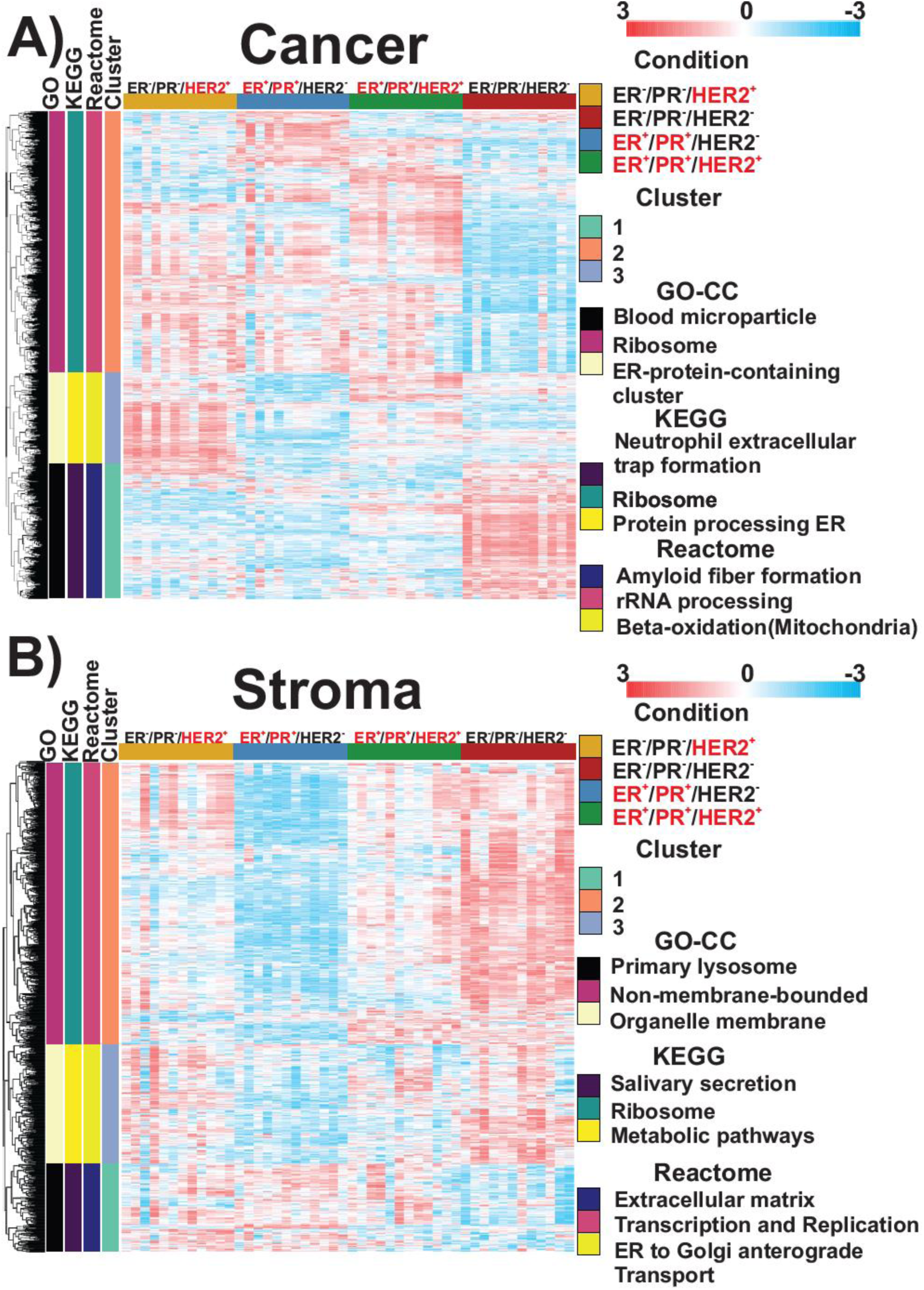
Spatial proteomic signatures across breast cancer molecular subtypes. **(A)** Hierarchical clustering of differentially expressed protein groups across ER⁻/PR⁻/HER2⁺, ER⁺/PR⁺/HER2⁺, ER⁺/PR⁺/HER2⁻, and triple-negative breast cancer subtypes, visualized independently for tumor (upper) and stromal (lower) compartments (n = 5 biological replicates; (two-way ANOVA; FDR 0.05).

**Supplementary Fig. 6.**
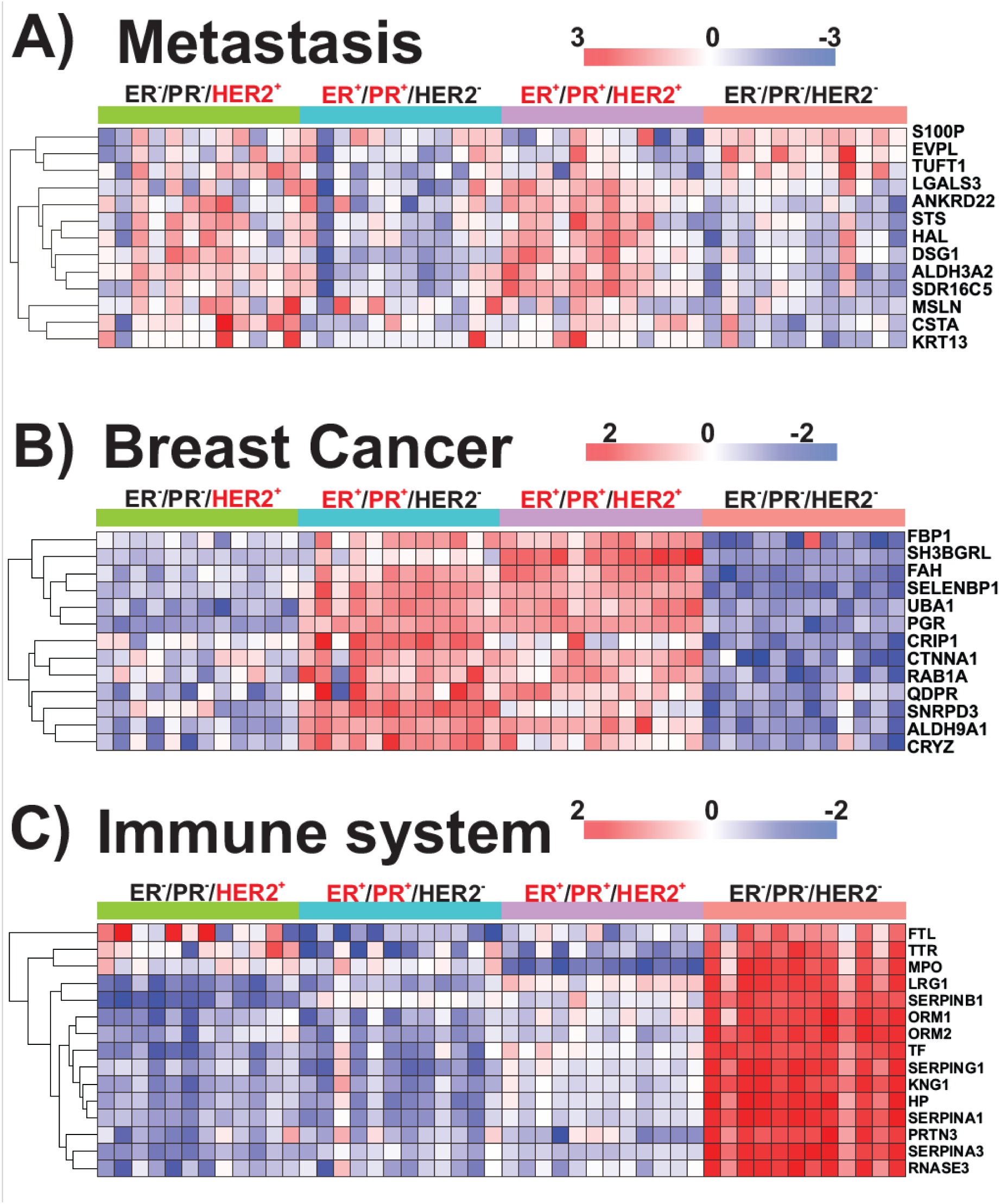
Spatial proteomic modular signatures across breast cancer subtypes. Hierarchical clustering of protein groups identified as differentially expressed by ANOVA across ER⁻/PR⁻/HER2⁺, ER⁺/PR⁺/HER2⁺, ER⁺/PR⁺/HER2⁻, and triple-negative breast cancer subtypes, shown for distinct functional modules: (A) metastasis-associated proteins, (B) breast cancer–associated signaling programs, and (C) immune-related signatures. Data are displayed separately by tumor subtype (n = 5 biological replicates; two-way ANOVA, FDR < 0.05).

**Supplementary Fig. 7.**
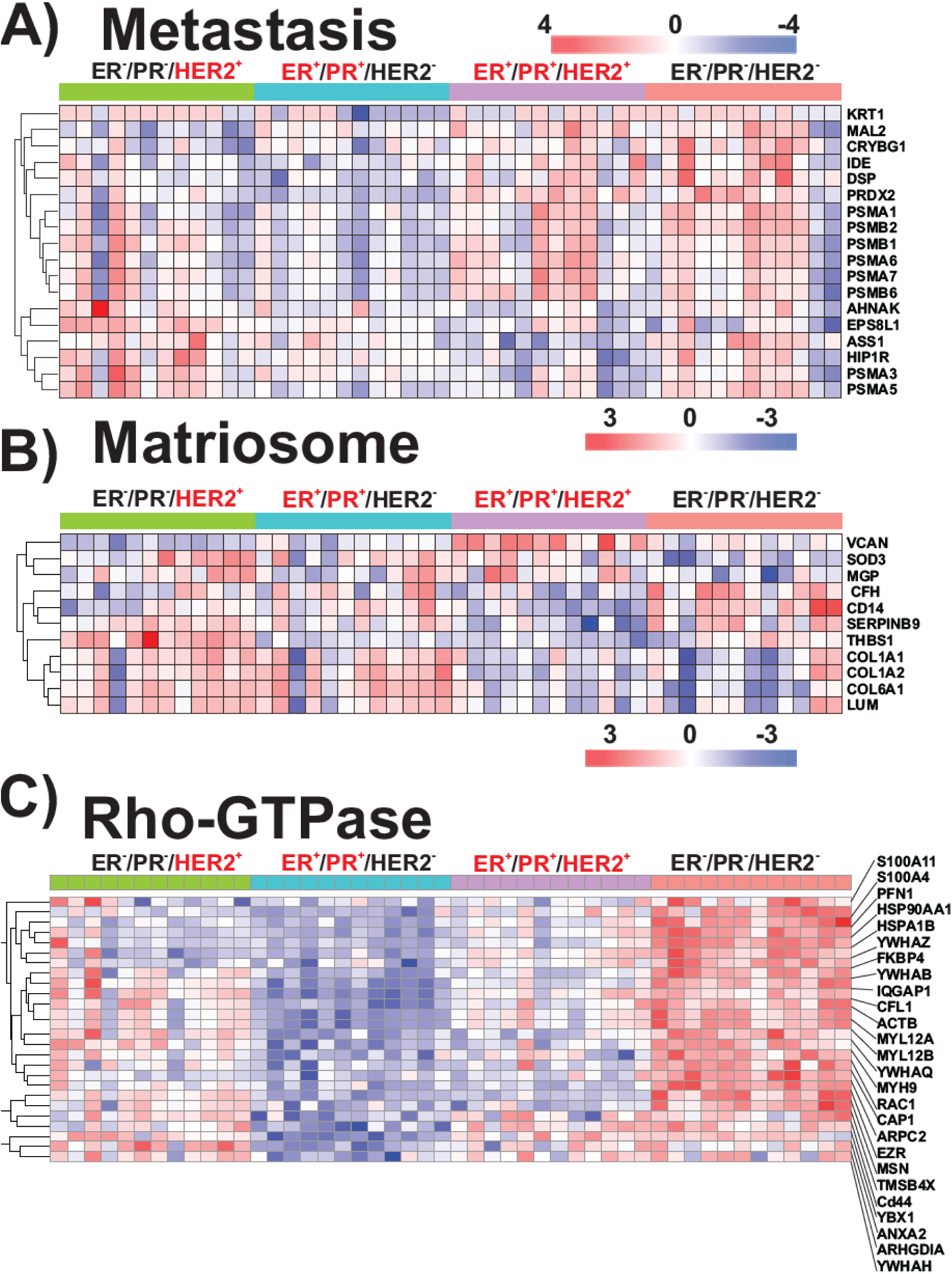
Spatial proteomic modular signatures across breast cancer stromal subtypes. Hierarchical clustering of protein groups identified as differentially expressed by ANOVA across ER⁻/PR⁻/HER2⁺, ER⁺/PR⁺/HER2⁺, ER⁺/PR⁺/HER2⁻, and triple-negative breast cancer stromal subtypes, shown for distinct functional modules: **(A)** metastasis-associated proteins, **(B)** Matriosome–associated signaling programs, and **(C)** Rho-GTPase-related signatures. Data are displayed separately by stromal tumor subtype (n = 5 biological replicates; two-way ANOVA, FDR < 0.05).

## References

1. Lundberg, E. & Borner, G. H. H. Spatial proteomics: a powerful discovery tool for cell biology. Nature Reviews Molecular Cell Biology vol. 20 285–302 Preprint at 10.1038/s41580-018-0094-y (2019).

2. Moffitt, J. R., Lundberg, E. & Heyn, H. The emerging landscape of spatial profiling technologies. Nature Reviews Genetics vol. 23 741–759 Preprint at 10.1038/s41576-022-00515-3 (2022).

3. Mund, A. et al. Deep Visual Proteomics defines single-cell identity and heterogeneity. Nat Biotechnol 40, 1231–1240 (2022).

4. Salgkamis, D. et al. Systematic review and feasibility study on pre-analytical factors and genomic analyses on archival formalin-fixed paraffin-embedded breast cancer tissue. Sci Rep 14, (2024).

5. Espina, V. et al. Laser-capture microdissection. Nat Protoc 1, 586–603 (2006).

6. Coscia, F. et al. A streamlined mass spectrometry–based proteomics workflow for large-scale FFPE tissue analysis. Journal of Pathology 251, 100–112 (2020).

7. Fowler, C. B., Waybright, T. J., Veenstra, T. D., OLeary, T. J. & Mason, J. T. Pressure-assisted protein extraction: A novel method for recovering proteins from archival tissue for proteomic analysis. J Proteome Res 11, 2602–2608 (2012).

8. Zhu, Y. et al. Nanodroplet processing platform for deep and quantitative proteome profiling of 10-100 mammalian cells. Nat Commun 9, (2018).

9. Makhmut, A., Qin, D., Hartlmayr, D., Seth, A. & Coscia, F. An Automated and Fast Sample Preparation Workflow for Laser Microdissection Guided Ultrasensitive Proteomics. Molecular and Cellular Proteomics 23, (2024).

10. Chen, H. et al. Routine Workflow of Spatial Proteomics on Micro-formalin-Fixed Paraffin-Embedded Tissues. Anal Chem 95, 16733–16743 (2023).

11. Kitata, R. B. et al. Robust collection and processing for label-free single voxel proteomics. Nature Communications 16, (2025).

12. Kwon, Y. et al. Hanging drop sample preparation improves sensitivity of spatial proteomics. Lab Chip 22, 2869–2877 (2022).

13. Chen, H. et al. Routine Workflow of Spatial Proteomics on Micro-formalin-Fixed Paraffin-Embedded Tissues. Anal Chem 95, 16733–16743 (2023).

14. Ye, Z. et al. One-Tip enables comprehensive proteome coverage in minimal cells and single zygotes. Nat Commun 15, (2024).

15. Guzman, U. H. et al. Ultra-fast label-free quantification and comprehensive proteome coverage with narrow-window data-independent acquisition. Nat Biotechnol 10.1038/s41587-023-02099-7 (2024) doi:10.1038/s41587-023-02099-7.

16. Ye, Z. et al. Enhanced sensitivity and scalability with a Chip-Tip workflow enables deep single-cell proteomics. Nat Methods 22, 499–509 (2025).

17. Roerden, M. & Spranger, S. Cancer immune evasion, immunoediting and intratumour heterogeneity. Nat Rev Immunol 25, 353–369 (2025).

18. Bennett, H. M., Stephenson, W., Rose, C. M. & Darmanis, S. Single-cell proteomics enabled by next-generation sequencing or mass spectrometry. Nat Methods 20, 363–374 (2023).

19. Petrosius, V. et al. Exploration of cell state heterogeneity using single-cell proteomics through sensitivity-tailored data-independent acquisition. Nat Commun 14, (2023).

20. Mund, A. et al. Deep Visual Proteomics defines single-cell identity and heterogeneity. Nat Biotechnol 40, 1231–1240 (2022).

21. Lundberg, E. & Borner, G. H. H. Spatial proteomics: a powerful discovery tool for cell biology. Nat Rev Mol Cell Biol 20, 285–302 (2019).

22. Lee, S. B., et al. Striking efficacy of a vaccine targeting TOP2A for triple-negative breast cancer immunoprevention. NPJ Precis Oncol 7, (2023).

23. Schroeder, J. et al. Investigating phenotypic plasticity due to toxicants with exposure disparities in primary human breast cells in vitro. Front Oncol 14, (2024).

24. Liu, N. Q. et al. Ferritin heavy chain in triple negative breast cancer: A favorable prognostic marker that relates to a cluster of differentiation 8 positive (CD8+) effector t-cell response. Molecular and Cellular Proteomics 13, 1814–1827 (2014).

25. Cheng, T. et al. CDKN2A-mediated molecular subtypes characterize the hallmarks of tumor microenvironment and guide precision medicine in triple-negative breast cancer. Front Immunol 13, (2022).

26. Chen, H. et al. KRT8 Serves as a Novel Biomarker for LUAD and Promotes Metastasis and EMT via NF-κB Signaling. Front Oncol 12, (2022).

27. Bodelon, C. et al. Mammary collagen architecture and its association with mammographic density and lesion severity among women undergoing image-guided breast biopsy. Breast Cancer Research 23, (2021).

28. Jansson, M. et al. Stromal Type I Collagen in Breast Cancer: Correlation to Prognostic Biomarkers and Prediction of Chemotherapy Response. Clin Breast Cancer 24, e360–e369.e4 (2024).

29. Hu, X., et al. Decorin-mediated suppression of tumorigenesis, invasion, and metastasis in inflammatory breast cancer. Commun Biol 4, (2021).

30. Lin, S. et al. Exploring the association of POSTN+ cancer-associated fibroblasts with triple-negative breast cancer. Int J Biol Macromol 268, (2024).

31. Strack, R. Subcellular spatial proteomics. Nat Methods 21, 2227 (2024).

32. Lv, Y. et al. Immune Cell Infiltration-Based Characterization of Triple-Negative Breast Cancer Predicts Prognosis and Chemotherapy Response Markers. Front Genet 12, (2021).

33. Sikandar, B., Qureshi, M. A., Naseem, S., Khan, S. & Mirza, T. Increased tumour infiltration of CD4+ and CD8+ T-lymphocytes in patients with triple negative breast cancer suggests susceptibility to immune therapy. Asian Pacific Journal of Cancer Prevention 18, 1827–1832 (2017).

34. Pereira, B. et al. The somatic mutation profiles of 2,433 breast cancers refines their genomic and transcriptomic landscapes. Nat Commun 7, (2016).

35. Kalluri, R. The biology and function of fibroblasts in cancer. Nature Reviews Cancer vol. 16 582–598 Preprint at 10.1038/nrc.2016.73 (2016).

36. Costa, A. et al. Fibroblast Heterogeneity and Immunosuppressive Environment in Human Breast Cancer. Cancer Cell 33, 463–479.e10 (2018).

37. Wu, S. Z. et al. Stromal cell diversity associated with immune evasion in human triple-negative breast cancer. EMBO J 39, (2020).

38. Maierthaler, M. et al. S100P and HYAL2 as prognostic markers for patients with triple-negative breast cancer. Exp Mol Pathol 99, 180–187 (2015).

39. Zhang, Y. S., Han, L., Yang, C., Liu, Y. J. & Zhang, X. M. Prognostic Value of LRG1 in Breast Cancer: A Retrospective Study. Oncol Res Treat 44, 36–41 (2021).

40. Chiao, C. C., et al. Prognostic and genomic analysis of proteasome 20s subunit alpha (PSMA) family members in breast cancer. Diagnostics 11, (2021).

41. Sundararajan, R. et al. Loss of correlated proteasomal subunit expression selectively promotes the 20SHigh state which underlies luminal breast tumorigenicity. Commun Biol 8, (2025).

42. Naba, A. et al. The matrisome: In silico definition and in vivo characterization by proteomics of normal and tumor extracellular matrices. Molecular and Cellular Proteomics 11, (2012).

43. Kaushik, S., Pickup, M. W. & Weaver, V. M. From transformation to metastasis: deconstructing the extracellular matrix in breast cancer. Cancer and Metastasis Reviews 35, 655–667 (2016).

44. Zhao, Y. et al. Stromal cells in the tumor microenvironment: accomplices of tumor progression? Cell Death and Disease vol. 14 Preprint at 10.1038/s41419-023-06110-6 (2023).

45. Kreuzaler, P. et al. Heterogeneity of Myc expression in breast cancer exposes pharmacological vulnerabilities revealed through executable mechanistic modeling. Proc Natl Acad Sci U S A 116, 22399–22408 (2019).

46. Horvath, P. & Coscia, F. Spatial proteomics in translational and clinical research. Molecular Systems Biology vol. 21 526–530 Preprint at 10.1038/s44320-025-00101-9 (2025).

47. Makhmut, A., Qin, D., Hartlmayr, D., Seth, A. & Coscia, F. An Automated and Fast Sample Preparation Workflow for Laser Microdissection Guided Ultrasensitive Proteomics. Molecular and Cellular Proteomics 23, 100750 (2024).

